# The effect of combined treatment of vitamin C and loperamide on intestinal sodium and potassium ion ATPase, alkaline phosphatase and lipid peroxidation on castor oil induced diarrheal rats

**DOI:** 10.1101/2020.11.13.380469

**Authors:** Aondowase Paul Iorhemba, Anthony Godswill Imolele

## Abstract

**Background:** Diarrhea is distinguished by prevalence of bowel movement accompanied by a loose consistency of stools, resulting from hyper peristalsis of the small intestine or colon, Diarrhea is a major challenge among infants and growing children. The study was carried out to assess the result of combined treatment of vitamin C and loperamide on intestinal Na^+^, K^+^ - ATPase, alkaline phosphatase, and lipid peroxidation in castor oil induced diarrheal wistar rats.

**Method:** A total of 18 wistar rats weighing 180-200g were randomly divided into 6 groups, (Group 1 Normal control no castor oil, no treatment administered, Group 2 Experimental control were given castor oil 3.0 ml/kg body weight with no treatment, Group 3 Standard control were given castor oil 3.0 ml/kg body weight + loperamide, Group 4 Treatment 1 were administered 3.0 ml/kg body weight + 25 mg/kg combined effect of vitamin C and loperamide, Group 5 Treatment 2 were administered 3.0 ml/kg body weight + 50 mg/kg combined effect of vitamin C and loperamide, and Group 6 were administered 3.0 ml/kg body weight + 100 mg/kg combined effect of vitamin C and loperamide) with 3 rats per group; the experiment lasted for 24 hours. The action of intestinal alkaline phosphatase, Na^+^, K^+^ - ATPase and malondialdehyde were determined.

**Result:** Descriptive statistical analysis was adopted using SPSS version 20. Combined effect of vitamin C and loperamide significantly (p<0.05) lowered the elevated levels of malondialdehyde caused by castor oil induced diarrhea; the Na^+^, K^+^ - ATPase intestinal activity treatment with both vitamin C and loperamide significantly elevated the activity of Na^+^, K^+^ ATPase when compared with the normal control, but both treatments (loperamide alone and vitamin C plus loperamide were not significantly different (p<0.05) to themselves. However, at 50 mg/kg body weight of combined effect of vitamin C and loperamide it showed significant difference in the action of intestinal alkaline phosphatase.

**Conclusion:** Findings of this study therefore, indicate that a combined effect of loperamide and vitamin C will be an effective therapeutic agent in the management of diarrhea by scavenging of free radicals generated in the cause of diarrheal to reduce lipid peroxidation. Therefore, combined effect of vitamin C and loperamide should be encouraged in the management of diarrhea. Further research should be directed towards assessing the therapeutic action of vitamin C only in the management of diarrhea.

## INTRODUCTION

### Background of the study

Diarrhea is an abdominal disorder, distinguished by a change in a usual bowel movement, which gives rise to a high volume of watery and persistence stools (Schiller *et al.,* 2017). An increase in the frequency of bowel flow causing discomfort and general body weakness (Wansi *et al*., 2014; Schiller *et al.,* 2017). It is generally accompanied by abdominal pain, stomach upset, urgency, rectal discomfort or pain, and fecal incontinence (Barr, 2014). It is not really a disease, and can be a manifestation of various conditions (Teferi *et al*., 2019). The epidemiology of diarrhea varies across low – middle – and high income countries (Wansi *et al*., 2014). The challenge of diarrheal is unarguably high in low – and middle income countries, as compared to high-income countries both in morbidity and mortality ratio (Teferi *et al.*, 2019). In developing countries, it is a major cause of death among children under 5 years and it is next to pneumonia in mortality ranking (Wansi *et al*., 2014). In Ethiopia, the mortality rate of diarrhea is almost half a million among children under 5 years and it is secondary to pneumonia (Gebru *et al*., 2014). Diarrhea accounts for almost 14% of hospital visits and approximately 16% hospital admissions (Umer *et al*., 2013). And despite the available standards of orthodox management and treatment of diarrhea, a high number of people in developing countries depend on herbs for the management and treatment of diarrhea (Umer *et al*., 2013). Currently the used medications in diarrhea treatment are essential in the management of diarrhea, but however, they are associated with adverse effects (FDA, 2016). A good example is loperamide and racecadotril used to treat and manage secretory diarrhea but they give rise to some symptoms such as bronchospasm, vomiting, and fever (Teferi *et al*., 2019). Cardiovascular complications are also attributed to excessive dose of loperamide (Teferi *et al*., 2019).

Vitamin C is a powerful antioxidant vitamin and its role played in various toxicity conditions in the body cannot be overemphasized because of its ability to scavenge free radicals that promotes toxicity (Danbature *et al.,* 2015). Several investigations have been carried out on physiological benefits of ascorbic acid and its application as therapeutic agents in various disease states (Hemila & Chalker, 2013).

A high number of major transporters has been linked in the pathogenesis of diarrheal diseases of which intestinal Na^+^/K^+^ ATPase is a very important one (Robinson & Flashner, 2008). Transporters are described as membrane proteins that regulate or mediate the movement of ions, small molecules and other macromolecules across the membrane (Ousingsawat *et al.*, 2011). The Na^+^/K^+^ ATPase enzyme is an example of the transporters which gives out 3 sodium ions for every 2 potassium ions absorbed by the cell, in order to maintain the cell biological integrity (Ousingsawat *et al.*, 2011). Intestinal epithelial cells regulate secretion and absorption of membrane electrolytes (Na^+^ K^+^ ATPase inclusive), which acts together to sustain fluid balance; however, this fluid balance is impaired during diarrhea (Robinson & Flashner, 2008).

Alkaline phosphatase located in the intestine is a major brush border enzyme that is produced throughout the abdominal region and discharged both to the bloodstream and the intestines (Krug *et al.,* 2014). lowered activity levels of intestinal alkaline phosphatase (IAP) could increase the risk of diarrheal (Krug *et al.,* 2014).

Lipid peroxidation is majorly destruction, due to the emergence of lipid peroxidation products which is seen during diarrhea leads to the spread of free radicals (Such as reactive oxygen and nitrogen species, ROS/RNS) (Pitocco *et al.,* 2010); that are responsible for a variety of degenerative process in the GIT infections like diarrhea (Pitocco *et al.,* 2010).

With the challenging health effect of diarrhea among children in developing countries this work therefore is designed to evaluate the result of combined treatment of vitamin C and loperamide on intestinal sodium and potassium ion ATPase, alkaline phosphatase and lipid peroxidation on castor oil and induced diarrheal wistar rats.

## MATERIALS AND METHODS

### Induction of Diarrhea

Diarrhea was induced to the rats with 3.0ml of Castor oil following the method reported by (Shah *et al.,* 2012). One hour later the rats were examined for the presence of characteristic diarrhea droppings, before treatment.

### Grouping of experimental animal

#### Animal grouping

The experimental wistar rats weighing 180-200g were randomly separated into six (6) groups, with 3 rats per group and were treated within a day and fed with commercial rat feed and water *ad libitum.*

| S/N | Groups | Treatments |
| --- | --- | --- |
| 1 | Normal control | No castor oil, no treatment |
| 2 | Experimental control | Castor oil 3.0ml/kg body weight No treatment |
| 3 | Standard control | Castor oil 3.0ml /kg body weight + loperamide |
| 4 | Treatment 1 | Castor oil 3.0ml/kg body weight + 25mg/kg body weight Vitamin C + loperamide |
| 5 | Treatment 2 | Castor oil 3.0ml /kg body weight + 50mg/kg body weight Vitamin C and loperamide |
| 6 | Treatment 3 | Castor oil 3.0 ml/kg body weight + 100 mg/kg body weight Vitamin C and loperamide |

### Sacrifice of the Animals

The animals were sacrificed one hour after the treatment.

### Collection of Samples

The intestines were dissected out carefully, following precautionary measures, the fluid was then emptied into a bottle for onward transportation to the laboratory for homogenization.

### Biochemical Assay

- **Alkaline phosphatase (ALP) activity:** Alkaline phosphatase activity was estimated using Agappe kit based on Schlebusch *et al.* (1974).
- **Lipid Peroxidation:**Malondialdehyde levels was estimated using the Thiobarbituric Acid reactions (Bhutia *et al*., 2011).
- **Na**^*+*^ - **K**^*+*^ **ATPase activity:** Na^+^ - K^+^ -ATPase activities was quantified by the method of Almansa *et al.* (2001).

### Statistical approach

Data obtained from this study was analyzed using descriptive statistics and presented as mean ± standard error of mean, using SPSS for windows version 20, difference between mean values were separated using the analysis of variance, followed by Duncan multiple test. The statistical significance was determined at *p* <0.05

## RESULTS

The result in Table 1 shows the concentration of malondialdehyde (MDA) in small intestinal homogenate of castor oil induced diarrheal rats. In assessing the lipid peroxidation castor oil induced diarrhea wistar rats showed an elevated level significantly (*p* <0.05) in MDA as compared to the normal control; which expresses the generation of free radicals (ROS) following diarrheal infection. However, following treatment at 25 and 50 mg/kg body weight of combined vitamin C and loperamide significantly (*p* <0.05) lowered the levels of MDA **(Table 1)**, but the combined effect of vitamin C and loperamide with a dose of 100 mg/kg body weight has shown to be more effective in the scavenging of free radicals generated during diarrheal infection as it significantly (*p* <0.05) lowered the MDA levels **(Table 1)**.

**Table 1:**
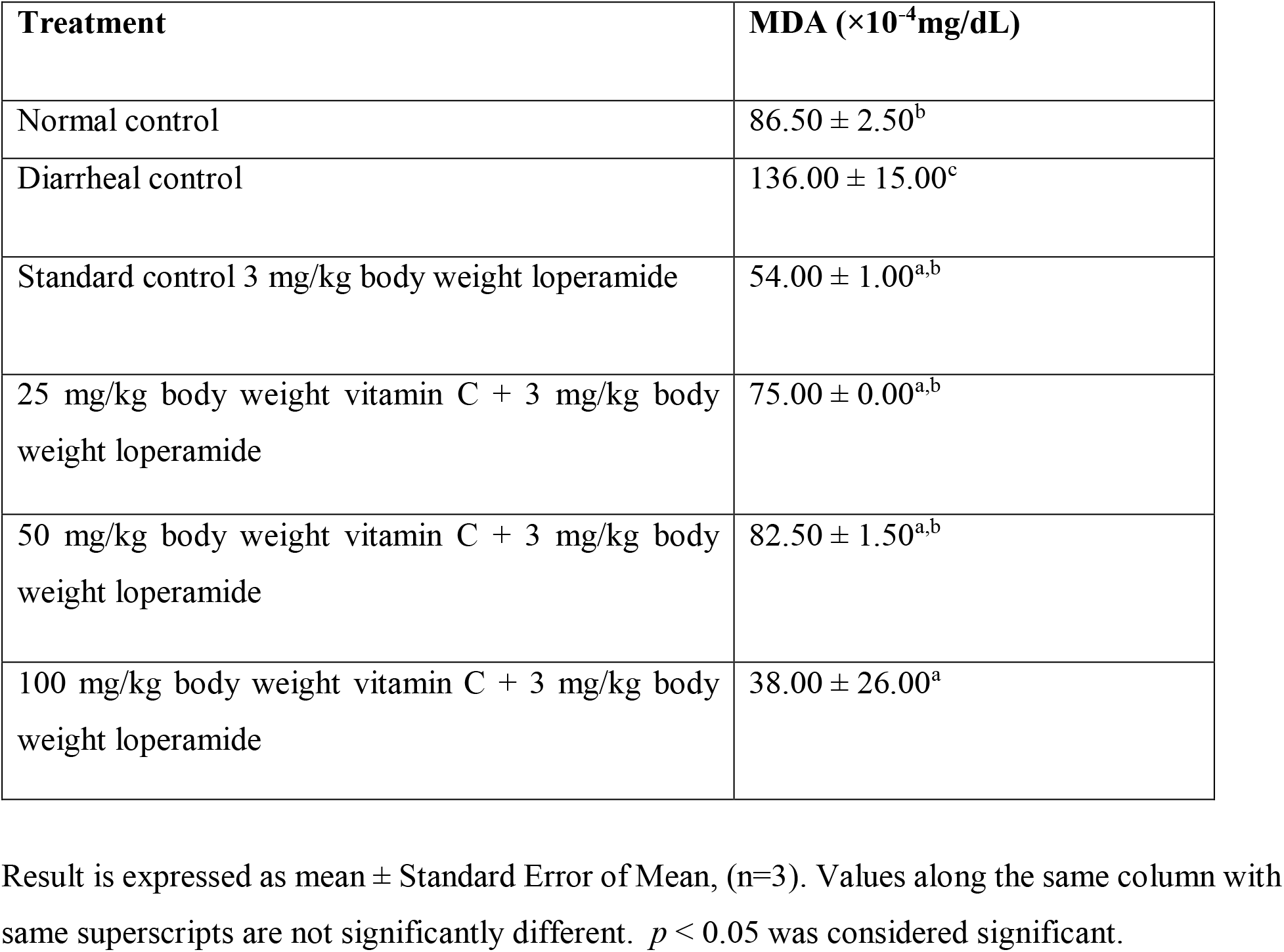
Concentration of Malondialdehyde (MDA) in small intestinal homogenate of castor oil induced diarrheal wistar rats.

| Treatment | MDA ($\times 10^{-4}$mg/dL) |
| --- | --- |
| Normal control | 86.50 ± 2.50<sup>b</sup> |
| Diarrheal control | 136.00 ± 15.00<sup>c</sup> |
| Standard control 3 mg/kg body weight loperamide | 54.00 ± 1.00<sup>a,b</sup> |
| 25 mg/kg body weight vitamin C + 3 mg/kg body weight loperamide | 75.00 ± 0.00<sup>a,b</sup> |
| 50 mg/kg body weight vitamin C + 3 mg/kg body weight loperamide | 82.50 ± 1.50<sup>a,b</sup> |
| 100 mg/kg body weight vitamin C + 3 mg/kg body weight loperamide | 38.00 ± 26.00<sup>a</sup> |

Result is expressed as mean ± Standard Error of Mean, (n=3). Values along the same column with same superscripts are not significantly different. $p < 0.05$ was considered significant.

The result above in Table 2 shows the activity of small intestinal alkaline phosphatase in castor oil induced diarrheal wistar rats. Castor oil induced diarrhea in wistar rats revealed a significant (*p* <0.05) decline in the activity of small intestinal alkaline phosphatase when compare with the normal control **(Table 2)**. However, there was significant increase in the activity of alkaline phosphatase following treatment with loperamide alone and as well as combined vitamin C and loperamide at 50 mg/kg body weight **(Table 2)**.

**Table 2:**
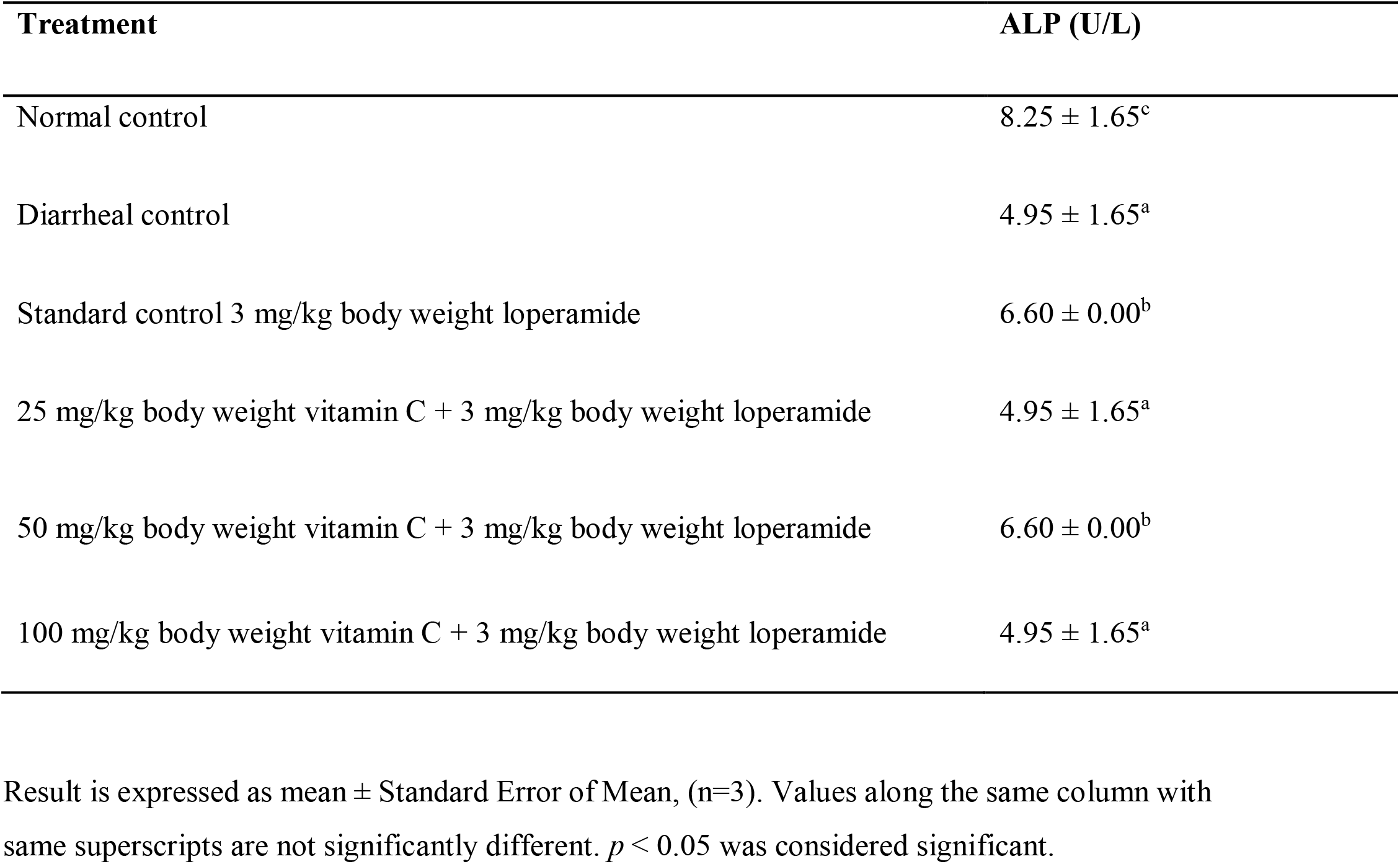
Activity of small intestinal alkaline phosphatase in castor oilduced diarrheal wistar rats.

The result above in Table 3 shows the activity of the small intestinal Na^*+*^/K^*+*^ -ATPase in castor oil induced diarrheal rats. Castor oil induced diarrhea in rats revealed a significant (*p* <0.05) decline in the activity levels of Na^*+*^/K^*+*^ - ATPase when compared with the normal control **(Table 3)**. However, upon treatment with loperamide and combined treatment of loperamide and vitamin C at varying dosages of 25, 50, and 100 mg/kg body weight revealed a significant (*p* <0.05) increase in the activity levels of Na^*+*^/K^*+*^ - ATPase enzyme when compared with diarrhea control group. **(Table 3)**.

**Table 3:** Activity of small intestinal sodium Na^*+*^/K^*+*^ - ATPase in castor oil induced diarrheal wistar rats.

| Treatment | $\text{Na}^+/\text{K}^+$ ATPase (U/L) |
| --- | --- |
| Normal control | $8.48 \pm 0.48^b$ |
| Diarrheal control | $5.34 \pm 1.13^a$ |
| Standard control 3 mg/kg body weight loperamide | $6.70 \pm 0.58^{a,b}$ |
| 25 mg/kg body weight vitamin C + 3 mg/kg body weight loperamide | $6.60 \pm 0.03^{a,b}$ |
| 50 mg/kg body weight vitamin C + 3 mg/kg body weight loperamide | $6.70 \pm 1.11^{a,b}$ |
| 100 mg/kg body weight vitamin C + 3 mg/kg body weight loperamide | $6.76 \pm 0.42^{a,b}$ |

Result is expressed as mean ± Standard Error of Mean, (n=3). Values along the same column with same superscripts are not significantly different. $p < 0.05$ was considered significant.

## Discussion

Diarrhea emerges from a disparity between the absorptive and secretory mechanisms in the abdominal tract, accompanied by hyper motility and leading to an excessive loss of fluid in the faeces (Gandhimathi *et al.,* 2009). In some diarrhea, the secretory components predominate while other diarrhea are characterized by hyper motility (Gandhimathi *et al.*, 2009). Castor oil, which is used to induce diarrhea is hydrolyzed in the upper small intestine to ricinoleic acid (Shah *et al.,* 2012), that produces inflammatory action on the mucosa of the intestines leading to the stimulation, formation and secretion of autacoids and other lipid compounds (Shah *et al.,* 2012). This however, cause the permeability of the mucosa cells to increase and alterations in electrolytes movement, which gives rise to hyper-secretory reaction (declining Na^*+*^/K^*+*^ absorption and reducing Na^*+*^/K^*+*^ - ATPase action in the small intestine and colon (Shah *et al.,* 2012); activation of adenylyl cyclase, platelet-activating factor and lately nitric oxide was discovered as a contributor to the diarrheal effect of castor oil.

In this study intestinal enzymes activity of alkaline phosphatase, Na^*+*^/K^*+*^ - ATPase and lipid peroxidation in castor oil induced diarrhea rats were determined and were compared with the values obtained from treated groups, with loperamide and combined effect of loperamide and vitamin C at varying dosages and as well as the normal control. Evidence showed that castor oil induced diarrheal significantly (*p* <0.05) increases the concentration levels of malondialdehyde which is a biomarker for the assessment of peroxidation of lipid and is line with the studies of (Shah *et al.,* 2012; Emmanuella, 2014) who described the generation of nitric oxide and other free radicals in castor oil induced diarrhea rats. However combined treatment of loperamide and vitamin C at 100 mg/kg body weight reduced significantly (*p* <0.05) the level of malondialdehyde, this could be attributed to the anti-oxidative capacity of vitamin C in scavenging free radicals which is consistence with the earlier reports of Abubakar *et al.* (2016) on antioxidative effect of vitamin C. Ascorbate provides a major function in the protection of plasma lipids from reactive oxygen species. Previous studies have reported vitamin C as a powerful antioxidant with the capability of scavenging free radicals thereby lowering the levels of peroxidation of lipids (Moncelli *et al.,* 2017; Mecan *et al.,* 2019).

Intestinal Alkaline phosphatase functions most importantly in the maintenance of the intestinal integrity by regulating bicarbonate secretion and duodenal surface pH, it functions also in the modulation of the intestinal long chain fatty acids (LCFA) absorption, and the detoxification of endotoxin lipopolysaccharide leading to intestinal and systematic anti-inflammatory effects (Bilski *et al.,* 2019). These actions of intestinal alkaline phosphatase are important in maintaining the biological integrity of the gut. In this study the intestinal alkaline phosphatase of diarrhea rats control decreased when compared to the normal control this is consistent with the findings of Bakare *et al.* (2010).

The Na^*+*^/K^*+*^ - ATPase, located in the basolateral membrane of the enterocytes functions as an important transporter of nutrients in the small intestine by moving K^*+*^ ions into and the Na^*+*^ out of the cell (Mabjeesh *et al.,* 2003) right within the enterocyte, homeostasis is maintained through active removal of sodium from the cell by the Na^*+*^, K^*+*^ - adenosine triphosphatase (ATPase) or sodium pump; because much of the movement of nutrient in the intestine is by sodium co-transporters, Na^*+*^, K^*+*^ -ATPase may be used to assess nutrients uptake (Mabjeesh *et al.,* 2003). In this study the activity of Na^*+*^, K^*+*^ - ATPase in castor oil induced diarrhea rats showed decreased level in the activity of Na^*+*^, K^*+*^ -ATPase, a number of physiological and biochemical mechanisms have been involved in the diarrhea effect of castor oil, these include: castor oil decreases fluid absorption, increases secretion in the small intestine and colon, and effects smooth muscle contractility in the intestine which is in line with the reports of (Meite *et al.,* 2009; Shah *et al.,* 2012; Emmanuella, 2014). Castor oil produce diarrhea effects by its functional components of recinoleic acid, inhibiting Na^*+*^, K^*+*^ - ATPase activity to reduce the physiological fluid absorption (Meite *et al.,* 2009). This could be attributed that both the combined treatment and the loperamide alone increased the activity. This suggest that the stimulatory effect may not be due to the addition of vitamin C.

## Conclusion

The findings of the study revealed that a combined effect of loperamide and vitamin C will be an effective therapeutic agent in the management of diarrhea by scavenging of free radicals generated in the cause of diarrheal infection and increasing ALP activity.

## Recommendations

Combined effect of vitamin C and loperamide should be encouraged in the management of diarrhea. Further research should be directed towards assessing the potency of vitamin C only in the management of diarrhea.

## Acknowledgement

We appreciate our mentors, Professor Eyong U. Eyong (University of Calabar), Dr. Oluwagbemiga Aina and Dr. Chika Onwuama, Nigerian Institute of Medical Research (NIMR) University of Lagos, Nigeria. Dr. Segun Fatumo, (Associate Professor) NCDE London School of Hygiene and Tropical Medicine, Group Leader the African Computational Genomics (TACG) Research Group, for their guidance throughout the course of this research. To Egbule Omoze Mercy and Okpaise Daniel; Molecular Genetics Scientist and Laboratory Manager (Covid), Department of Molecular Diagnostics 54gene, for their great encouragement and advice.

